# Spectral estimation at the edge

**DOI:** 10.1101/2024.10.02.616083

**Authors:** Shivangi Patel, Eleni Psarou, Gregor Mönke, Pascal Fries

## Abstract

Cognitive functions depend on neuronal communication, which is subserved by the synchronization of neuronal rhythms. Rhythms are characterized by their frequency, power and phase. If the phase of a rhythm just preceding an input is predictive of the neuronal or behavioral response to the input, this provides strong evidence for a functional role of the rhythm. Yet, this requires estimating the phase of a rhythm at the edge of the epoch. This is challenging, because any phase estimation that is spectrally specific requires a finite window length often combined with tapers that de-emphasize the signal close to the edge. To overcome this, we propose a method that builds on previously described approaches based on autoregressive modeling of the data and corresponding extrapolation beyond the edge. In contrast to related previous approaches, the modeling is based on the broadband signals, avoiding filtering-related group delays, and the extrapolation is performed multiple times, allowing averaging and thereby the reduction of extrapolation noise. The new method provided more accurate phase estimation at the edge for most simulated datasets, and for an empirical dataset from awake macaque area V4. We propose that the enhanced phase estimation accuracy at the edge might help to investigate the functional roles of brain rhythms and potentially also to improve phase-specific stimulation for clinical applications.

## 1. Introduction

Cognitive functions have often been linked to brain rhythms. Rhythms can be characterized by their frequency, power and phase. These parameters together capture the rhythmic state of a neuronal system. This state might influence how the system responds to input. To investigate whether the system’s input-output characteristic depends on the rhythm’s state, one needs to estimate the rhythm’s state prior to the input, to correlate it with the subsequent system output.

This analysis can be performed for each of the rhythm’s parameters, and numerous studies have linked neuronal or behavioral responses to the frequency of brain rhythms, their power or their phase (Busch et al., 2009; Harris et al., 2018; van Vugt et al., 2018). We argue that an observed relation between a rhythm’s phase and a system’s response makes a particularly strong case for a functional role of the rhythm. This is, because the phase values undergo a full rotation during each cycle. If these rapidly rotating values are predictive of a system’s response, then any confounding variable would need to show correlated rotations, synchronized to the rhythm’s phase. While this is in principle conceivable, it is much more parsimonious to conclude that the function is due to the phase, and thereby due to the rhythm, itself. The cognitive function of a brain rhythm’s phase has indeed been demonstrated in several studies. For example, a seminal study showed in human participants that neuronal and behavioral responses to a visual probe stimulus partly depend on the phase of a frontal theta rhythm (Busch et al., 2009). A later study showed in awake macaque monkeys that both neuronal and behavioral responses to a stimulus change partly depend on the phase of the stimulus-induced visual cortical gamma rhythm (Ni et al., 2016).

The relation between a rhythm’s phase at a given time point and a subsequent neuronal or behavioral response can only be investigated when the phase estimation is not confounded by the response. That is, the response needs to be excluded from the epoch that is used for the estimation of the phase. Note that the estimation of phase for a specific frequency or frequency band always requires a signal of some length of time, that is, a signal epoch. This signal epoch is typically multiplied with a taper in order to avoid aliasing, and then Fourier transformed. The taper approaches a value of one in its center and zero on both ends, and thereby emphasizes the center of the epoch and de-emphasizes both ends of the epoch. However, the right end of the epoch is closest to the response that is supposed to be predicted by the preceding phase. Therefore, this part of the epoch should not be de-emphasized. This is particularly relevant, because physiologically occurring rhythms often show substantial phase drift, such that the phase at the end of the epoch can have drifted substantially from the phase predicted on the basis of the center of the epoch. Therefore, we developed methods aimed at optimizing the estimation of spectral phase at the edge of a signal epoch.

Several methods have been proposed for phase estimation at the edge of an epoch. We consider three classes of methods, based on [1] asymmetric tapers, [2] endpoint-corrected Hilbert transform and [3] signal extrapolation. In all methods, the signal epoch used for spectral estimation ends at the ‘critical time’, such that the epoch excludes the response (or even the stimulus or probe that evokes the response). The asymmetric-taper-based methods involve designing frequency-dependent tapers, whose peak is shifted towards the end, in order to de-emphasize the center and emphasize the end (Mitchell et al., 2007). One recently described method is based on an endpoint-corrected Hilbert transform, and uses filtering in the frequency domain to reduce the so-called Gibbs phenomenon (Schreglmann et al., 2021). The extrapolation-based methods use regular, symmetric tapers, centered on the critical time to give maximal emphasis to the end of the signal epoch; in order to enable this, they extrapolate the signal beyond the critical time; this in turn is typically preceded by band-pass filtering of the signal to simplify extrapolation (Chen et al., 2013; Ni et al., 2016; Blackwood et al., 2018; Zrenner et al., 2020; Wodeyar et al., 2021; Wodeyar et al., 2023). However, the band-pass filtering can introduce substantial group delays that vary across frequencies. Such delays might move the effective time of the critical time in a frequency-dependent way that could significantly complicate the analysis. To address this issue, we propose a method based on autoregressive (AR) extrapolation, which avoids band-pass filtering by using the broadband signal, and which includes a way to average over multiple realizations of the extrapolation.

We compare the spectral phase estimation accuracy of our method to previously published methods by applying them to both simulated and empirical data. The simulated data is constructed by simulating oscillators with phase diffusion and adding pink noise to approximate neuronal signals. In these simulated data, the phase of the oscillators at any time is known by construction. We investigate how phase diffusion and noise strength influence the accuracy of phase estimation. At last we compare the estimation performance on empirical data without a response (or stimulus or probe), such that the phase at a ‘surrogate critical time’ could be estimated with the standard Hann taper method. For the empirical and most of the simulated data, the method proposed here obtained a higher spectral phase estimation accuracy than the previously published methods against which we compared.

## 2. Methods

### 2.1. Description

In short, our approach is based on the previously described extrapolation methods in which the signal epoch ends at the critical time. An AR model is fitted to the broadband signal before the critical time and used to extrapolate the broadband signal beyond the critical time. Subsequently, a Hann taper is centered at the critical time, and a Fourier transform is used for calculating the phase. To the best of our knowledge, our method differs from previous methods in the way in which the AR model is fit and used (Chen et al., 2013; Ni et al., 2016). The following paragraphs describe the method in more detail.

The rationale for our approach is two-fold: [1] We need to discard the signal after the critical time, such that it cannot confound our spectral estimate. The spectral estimate is supposed to be used to potentially explain the response to the event happening at the critical time, and therefore the spectral estimation must exclude the response, and correspondingly the signal must end at the critical time. [2] The taper should ideally be centered on the critical time, such that the signal around the critical time is least reduced by the taper function and maximally contributes to the spectral estimate. To achieve both [1] and [2], we need to use the signal only up to the critical time, then extrapolate it beyond the critical time, subsequently apply the taper centered to the critical time and apply the Fourier transform.

For the extrapolation, we used an autoregressive (AR) model. The AR model attempts to describe the signal at a given time by the signal at previous times multiplied with the respective model coefficients. Thereby, the model coefficients capture regularities in the signal. Once the coefficients have been fitted to the signal, they can be used with the signal of a given trial to extrapolate that trial’s signal. This extrapolation is expected to maintain the signal’s overall regularities, i.e., the extrapolated signal has similar power spectra and smooth phase progression from the signal that was extrapolated.

The AR model was fitted to the signal preceding the critical time. The signal was cut into epochs that were end-aligned to the critical time, and that had a length as specified below for the different analyses. Each epoch was detrended and subsequently z-scored by subtracting its mean and dividing by its SD. In our specific implementation of the AR fit, we chose to fit one AR model to all epochs, yet the AR model fit could also have been performed per epoch.

Before the AR model can be fitted, its model order needs to be determined, and this is explained further below. The AR model fit itself was implemented using a collection of MATLAB modules called ARfit (Neumaier and Schneider, 2001; Schneider and Neumaier, 2001). The modules in ARfit use an *n*-variate autoregressive model of order *p*

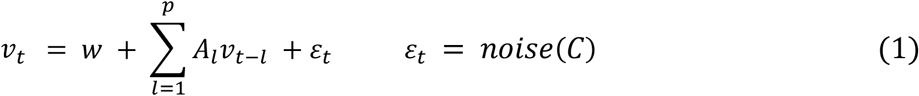

This is a model for stationary time series with vectors *v*_*t*_ such that they have been recorded at equally spaced intervals *t*. These vectors are of *n* dimensions, with *n* being different trials in our case. The order *p* determines the number of past vectors, in our case time steps, that the model takes into account. *A*_1_, *A*_2_, …….., *A*_*p*_ ∈ *R*^*n*×*n*^ are the coefficient matrices of the AR model. *ε*_*t*_ = *noise*(*C*) are uncorrelated random vectors with zero mean and covariance matrix *C* ∈ *R*^*n*×*n*^. *w* ∈ *R* ^*n*^ is an intercept vector that allows the mean of the time series to be nonzero.

The ARfit toolbox has a function called *arfit* that lets us calculate the coefficient matrices *A*_1_,*A*_2_,………,*A*_*p*_, intercept vector *w* and covariance matrix *C* given the model order.

The appropriate model order can be determined using different criteria like the Bayesian Information Criterion (BIC) or the Akaike Information Criterion (AIC), given the data and a specification of the minimum and maximum model order considered. We used the BIC and specified model orders to be between 1 and 300. The ARfit function used for order calculation is called *arord*. Accuracy of the AR extrapolation can be checked by the *arres* function which plots the autocorrelation of the residuals. For the model to be adequate, residuals have to be uncorrelated. If 95% of the autocorrelations (for lag > 0) lie within the confidence limits for the autocorrelations of an IID (independent and identically distributed random variable) process of the same length as the time series of residuals, then we consider the model order to be accurate.

We performed additional tests to assess the quality of the extrapolated signals. First, we plotted the signal up to the critical time along with the extrapolated signal after the critical time, to visually inspect whether the signal progresses smoothly around the critical time. Second, we plotted the power spectra of the signal up to the critical time and of the extrapolated signal, to visually inspect whether they are similar. This was performed for numerous trials. We found that the selected model order (specified later for the different signal types) was appropriate. Note that the appropriate model order also depends on the sampling rate of the signal. The sampling rate in our case was 1000 Hz. Several previous studies using AR models first decimated the sampling rate, which typically results in a lower required model order. We refrained from such decimation, because it entails low-pass filtering with the corresponding side effects.

After obtaining the appropriate model order, one AR model was fitted to the broadband signal epochs of all trials, using the selected model order. The fitted AR model was used, together with each individual epoch to extrapolate each epoch (using the ARfit toolbox). We refer to the individual epochs of each trial end-aligned to the critical time as the original epochs. We refer to an AR-based extrapolation of the original epoch as an extrapolated epoch. We refer to an original epoch concatenated with an extrapolated epoch as a compound epoch. The compound epoch was spectrally analyzed for frequencies from 2 to 60 Hz, in steps of 2 Hz. For each frequency, a Hann taper of three-cycle length, centered at the critical time, was multiplied with the compound epoch, and the Fourier transform was applied (using the ‘mtmconvol’ method of the ft_freqanalysis function of the FieldTrip toolbox (Oostenveld et al., 2011)).

We reasoned that one single extrapolation produces a noisy estimate, because half of the Fourier transformed signal (the extrapolated epoch) is based not only on the original epoch, but also on the white noise required to drive the AR model. As the original epoch is extrapolated time bin by time bin, the extrapolated signal is less and less defined by the original epoch. To minimize the influence of this randomness in the extrapolated epoch, we devised a novel approach to average over multiple extrapolations. For each original epoch, we ran the AR-based extrapolation 100 times and thereby obtained 100 compound epochs. If we had averaged those 100 compound epochs in the time domain, the extrapolated epochs would have increasingly cancelled each other (with this effect increasing with distance from the critical time). To avoid this cancellation, we first Fourier transformed the compound epochs. For each compound epoch, and for each frequency, the Fourier transform provides one complex vector, with an amplitude and a phase. The core methodological innovation here is that we determined the average phase and the average amplitude separately, and subsequently combined the average phase with the average amplitude into a new complex vector. This approach avoids the abovementioned cancellation, yet achieves the elimination of randomness.

In formal terms: Let *F*_*i*_ be the array of Fourier transforms of all 100 compound epochs derived from one single original epoch. Then the average phase is calculated as

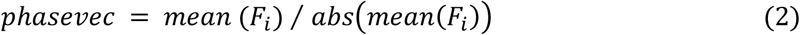

The average amplitude is calculated as

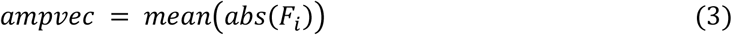

The average Fourier transform is then composed as

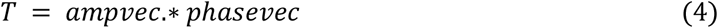

To find the final phase at a given frequency at each trial, we obtain the angle of this transform (*T*)

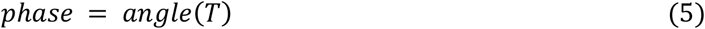

Note that the phase obtained in equation (2) could also have been used directly; we recomposed the phase with the amplitude to obtain the average Fourier transform, because it can be useful for many purposes.

We refer to this method as the AR-Fourier method from here on. A schematic for the method is described in Figure 1.

**Figure 1.**
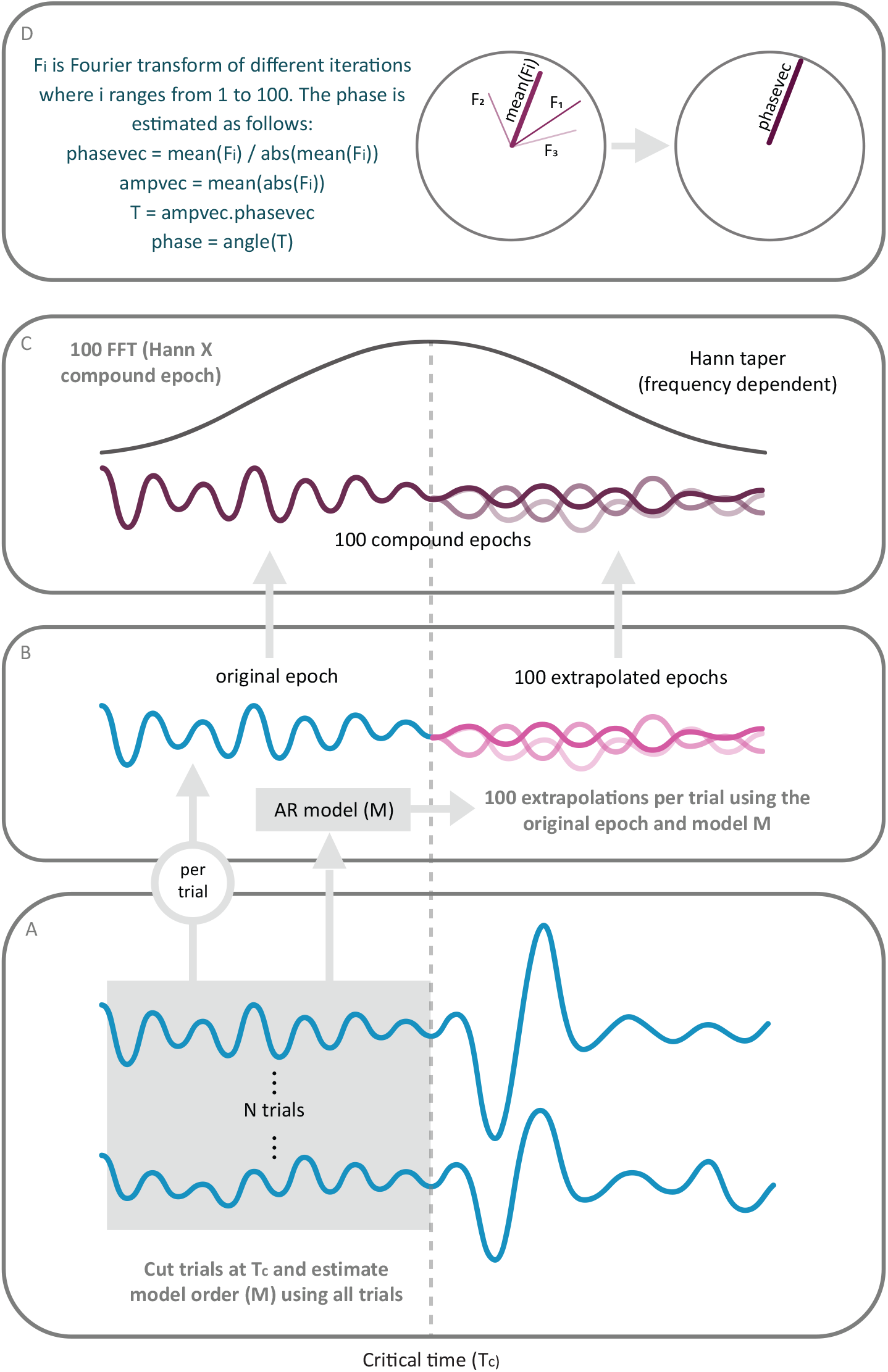
Schematic for the AR Fourier method. (A) N trials of the signal are shown. T_c_ is the critical time. All trials of the original signal before T_c_ are used to estimate the AR model. (B) The AR model is used together with each original epoch to generate 100 extrapolated epochs per original epoch. (C) Each extrapolated epoch combined with the corresponding original epoch constitutes a compound epoch. Each compound epoch is combined with a frequency-dependent Hann taper and subsequently Fourier transformed. (D) The 100 compound epochs derived from each original each lead to 100 Fourier transforms. For each Fourier transform, the figure shows the Fourier vector (as thin line) for one example frequency, illustrating the frequency’s amplitude and phase. The vectors of the 100 compound epochs are complex averaged. The phase of the average is the result of the AR-Fourier method.

### 2.2. Testing Method

#### 2.2.1. Using simulated data

##### 2.2.1.1. Phase Diffusion Model

In order to test our method, we created simulated data that allowed us to know the true instantaneous phase at certain frequencies.

The temporal derivative of the phase can be written as follows

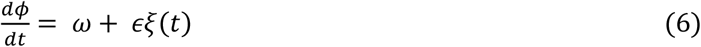

Here, *ω* is the constant angular frequency and *ξ*(*t*) is Gaussian white noise at time t with diffusion constant *ξ*.

For the integrated process, the phase at time *t* can be written as

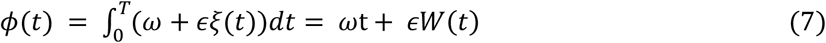

Where *W*(*t*) is the Wiener Process describing diffusive phase dynamics, and *ωt* is the phase velocity or the linear drift term. In this form, the degree of phase diffusion around the linear phase propagation is specified by the diffusion constant *ϵ* (Goldobin et al., 2003).

Numerical solutions of equation (7) give realizations of the unwrapped phase, which we subsequently transform to a time series by taking the cosine to obtain the scalar signal *x*(*t*)

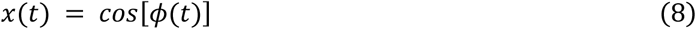

Note that the time-averaged angular frequency remains omega

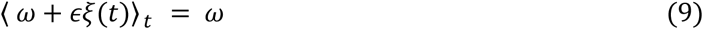

because the white noise *ξ*(*t*) is a zero-mean process. This gives us a concentrated spectral peak around *f* =*ω/*2*π* even with phase diffusion, when *ϵ* << *ω*

When we use white noise in the phase diffusion model, the resulting signal (*x*(*t*) in equation (8)) has some high frequency components which might make phase comparisons difficult. Therefore, in the model, we used a moving average filter (Gaussian window) on the white noise in equation (6) to reduce the high frequency component, while maintaining the overall strength of phase diffusion. In order to compare models with and without this filtering, we compared their average autocorrelation and time-dependent coherence and found them largely conserved after the filtering.

##### 2.2.1.2. Model for simulated data

The simulated signal was the sum of two frequency components, the target component for which we intended to estimate the phase, and the distractor component that was intended to mimic concurrent components in empirical data. The two signals had two different average angular frequencies, generated according to equation 8, and an AR(1) process that was used as a proxy for 1/f noise:

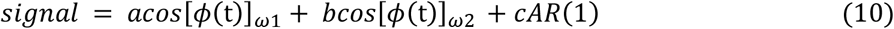

Here, *ω*1 and *ω*2 are different frequencies at which the true phase is known by construction. The true phase of the signal is [*ϕ*(*t*)]_*ω*1_ and [*ϕ*(*t*)]_*ω*2_ for the frequencies *ω*1 and *ω*2, respectively. Here, the parameter *c* was used to scale the strength of the AR1 process.

##### 2.2.1.3. Parameters of simulated data

We chose the frequencies *ω*1 and *ω*2 to be 4 Hz and 6 Hz, respectively. Five datasets were created by varying the values of *c* and *c* to obtain different proportions of these two frequency components. The value of *c* was set to be 1, and the value of *c* was varied to be 0.2, 0.4, 0.6, 0.8 and 1. This was done to investigate how the accuracy of the later phase estimation for the target component (*ω*1) was affected by varying proportions of the distractor component (*ω*2).

Furthermore, for each of these five datasets, pink noise strength could assume the values [0.1, 0.2, 0.3 … 0.9], and phase diffusion could assume the values [0.001, 0.051, 0.101, 0.151, 0.201, 0.251, 0.301, 0.351, 0.401, 0.451]. Phase diffusion was capped at 0.451, because above that value, the diffusion part of the phase is much stronger than the linear drift due to the underlying oscillation and therefore, the concept of phase does not anymore apply. Thus, for each of the five parameter values for *ω*2, there were a total of 90 combinations of the parameters for pink noise and phase diffusion, resulting in a total of 450 parameter sets. For each parameter set, a total of 500 trials were simulated, and each trial was 10 s long.

##### 2.2.1.4. Application of AR-Fourier Method on Simulated Data

Each 10 s trial was cut in half, and phase estimation was performed at the right-side edge of these epochs for the 4 Hz frequency. Model order was determined separately for each dataset, combining all respective 5 s long epochs, and the resulting value for the model order varied depending on the complexity of the simulated signal. The phase estimation followed the method description in section 2.1.

#### 2.2.2. Using Empirical data

##### 2.2.2.1. Data description

We tested our method also on empirical data, in particular local field potentials (LFP) recorded in area V4 of one awake macaque monkey. The monkey was implanted with electrode arrays (‘Utah arrays’) in areas V1 and V4 of the left hemisphere; only V4 data were used here. The V4 recording sites had receptive field (RFs) in the lower right quadrant. The empirical dataset was from one recording session with 500 trials, in which signals were obtained from 62 recordings sites. The signals were acquired with Tucker Davis Technologies systems (TDT, Alachua, Florida, United States). They were digitized at 24414.0625 Hz using a TDT PZ2 preamplifier. Offline, the signals were downsampled to 1000 Hz. The LFP was obtained by first upsampling to 3,125,000 Hz, then downsampling to 25,000 Hz, subsequently applying an IIR filter with a stop-band at 500 Hz and finally downsampling to 1000 Hz.

Each trial started with the presentation of a fixation point and a rectangular bar in the lower right quadrant, overlapping the V4 RFs during fixation. Once fixation was attained, a random time from 1 to 3 s elapsed until a small probe stimulus (a Gaussian contrast decrement) was presented at a random position on the bar. The aim of the experiment was to investigate the response to the probe stimulus as a function of the spectral phase right before the probe presentation. The optimal estimation of this spectral phase was the motivation for the development of the method presented here. When we will, in a later study, use our method for estimating the pre-probe phase, we will align the epochs to the probe presentation, and exclude the data affected by the probe and replace it by the AR-extrapolated data. In the present study, we aim at presenting and testing the method, and therefore, we needed to follow a different approach. We needed to compare the phase estimate by our method to the ground-truth phase estimate, referred to as ‘true phase’ (acknowledging that there is not literally a true phase for empirical data). The true phase at time *t* for the empirical data is defined here as the estimate based on a Hann-tapered Fourier transform centered on *t*. We used a 1-s epoch end-aligned to the probe onset, completely devoid of responses to the probe. The true phase is then the estimate of the spectral phase at the center of this 1-s epoch. This center was defined as the critical time (for the analysis presented here), at which we intend to estimate the phase without the influence of any data recorded after this critical time.

##### 2.2.2.2. Application of AR-Fourier Method on Empirical Data

We first used the FOOOF method (Donoghue et al., 2020; Gerster et al., 2022) to estimate the major spectral components, which revealed a dominant component at approximately 10 Hz. We therefore evaluated the quality of phase estimation at 10 Hz. As described in the previous paragraph, we used 1 s epochs end-aligned to stimulus onset, we used a Hann taper for these complete epochs to estimate the true phase, and we used the first 0.5 s to estimate the phase at the critical time in the center of the 1 s epoch. A model order of 50 was used for extrapolation, and the procedure described in section 2.1 was followed.

### 2.3. Comparison to previously established methods

#### 2.3.1. Previously Established Methods

We compared the performance of the AR-Fourier based method to previously published methods that either explicitly aimed at spectral estimation at the edge or that appeared like strong candidates for good spectral estimation at the edge. These previously described methods can be broadly classified in three groups, namely extrapolation-based methods, taper-based methods and endpoint-corrected Hilbert transform method.

##### 2.3.1.1. Extrapolation based methods

This class of methods relies on band-pass filtering the data in the required frequency range, extrapolating the data and taking the Hilbert transform for phase estimation. The extrapolation can be done using two different types of models, namely autoregressive models and state-space models.

The general rationale of autoregressive (AR) based extrapolation has been described above. Previous studies using AR-based extrapolation differ from our approach in [1] using band-pass filtered data, [2] using a single extrapolation. AR-based extrapolation has been implemented by several previous papers (Chen et al., 2013; Blackwood et al., 2018; Zrenner et al., 2020). We will refer to them by the name of their first author and their publication year as Chen2013 (Chen et al., 2013), Blackwood2018 (Blackwood et al., 2018) and Zrenner2020 (Zrenner et al., 2020). We compare our method to Blackwood2018 and Zrenner2020, because they have been shown to outperform Chen2013. Note that Wodeyar et al. (2021), referred to here as Wodeyar2021, has already compared Blackwood2018 and Zrenner2020 to each other and to their own method, and here we have used the implementations of these three methods published with Wodeyar2021.

Wodeyar2021 proposes extrapolation by means of a state-space model. The signal is modelled as a state-space oscillator or a combination of oscillators driven by noise. These models are used to describe and predict the behavior of a system over time. They consist of two equations: the state equation tries to model the hidden, underlying dynamics of the system we are interested in estimating. And the observation equation converts the hidden dynamical state (that is phase in this case) into the observed signal that we can measure or observe. To make the best prediction for the hidden state, Wodeyar2021 uses a Kalman filter. It compares the initial prediction of the observation with the actual observed value and updates the estimate in real-time.

##### 2.3.1.2. Taper-based methods

Hann tapers are excellent tapers for spectral estimation when they can be centered at the time for which the spectral estimate is intended. However, for spectral estimation at the edge, Hann tapers suffer from the fact that they strongly taper down the signal close to the epoch edges, such that the estimate is dominated by the center of the taper, respectively the center of the epoch. One might consider to linearly extrapolate this phase estimate from the epoch’s center to the edge. However, from the epoch center to the right epoch edge, there can be substantial phase diffusion. This phase diffusion moves the true phase at the right edge substantially away from the phase estimated at the center. Thus, using a Hann-taper-based approach for spectral estimation at the edge does not appear promising. Nevertheless, as Hann-taper-based estimation is probably the most commonly used method for general spectral estimation, we included this method here for comparison and refer to it as Hann. For each frequency, the Hann taper had a length of two cycles and was end-aligned to the critical time. Note that the AR-Fourier method used three cycles centered to the critical time, corresponding to 1.5 cycles before the critical time. We intended to design the Hann method as comparable as possible, and therefore opted for two cycles. The Hann method was implemented in FieldTrip, using freqanalysis with the mtmconvol option.

Note that the same considerations as for the Hann taper apply for any symmetric taper, like the Gaussian taper used in Morlet wavelets. Several previous studies have used wavelets for spectral estimation at the edge (Busch and VanRullen, 2010) and have found strong and convincing relations between the phase before the critical time and neuronal and/or behavioral responses after the critical time. The fact that they found this despite using a relatively insensitive method for spectral estimation at the edge does not call their findings into question, but rather suggests that they underestimated the strength of the described effect.

Another frequently investigated approach for spectral estimation at the edge of an epoch involves the utilization of asymmetric tapers. These tapers are designed to accentuate the latter portion of the epoch more than conventional tapers. Since the aim is to give greater weight to the signal closer to the epoch’s edge, employing these tapers, especially when they are aligned with the epoch’s end, can enhance phase estimation accuracy. In our study, we utilized two types of asymmetric tapers. The first one is the Alpha taper that we refer to as Mitchell2007 (Mitchell et al., 2007). The second one is the asymmetric Hann taper that we refer to as Asbai2014 (Asbai et al., 2014). Note that Asbai2014 is not focused on this aspect, but generally explains how to introduce asymmetry in a taper, which we apply here to the Hann taper. The method of Mitchell2007 is available as an option in FieldTrip and requires at least 5 cycles of each estimated frequency. We used this FieldTrip option with 5 cycles, end-aligned to the critical time. This entailed that we could use it only on the simulated data, but not on the empirical data, for which not enough data before the critical time was available. The method of Asbai2014 was implemented by introducing a custom asymmetric Hann taper function into the FieldTrip toolbox. This involved using two cycles of the Hann taper, end-aligned to the critical time, resulting in varying taper lengths for different frequencies. Both Mitchell2007 and Asbai2014 were implemented using the ‘mtmconvol’ method within the ft_freqanalysis function of the FieldTrip toolbox.

##### 2.3.1.3. Endpoint corrected Hilbert transform method

In addition to the Hann method and the asymmetric methods, we also included phase estimation comparisons using the endpoint-corrected Hilbert transform (ecHT) method referred here as Schreglmann2021 (Schreglmann et al., 2021). This method essentially uses the Hilbert transform to find the phase and employs a causal bandpass filter on the frequency domain representation of the signal to mitigate the Gibbs phenomenon. The implementation of the Schreglmann2021 method was based on the code provided in the supplementary material of the corresponding paper (Schreglmann et al., 2021).

##### 2.3.1.4. General remarks pertaining to all methods

Some of these methods have been described for data epochs substantially longer than the few cycles required for the AR-Fourier method. This holds e.g. for the state-space-based method of Wodeyar2021. We had 5 s epochs available for our simulated data and therefore provided them for all tested methods. For the empirical data, we had merely 1 s in total, and 0.5 s before the critical time (see above). Therefore, we adapted the methods that were previously described for longer epochs to apply them to these shorter empirical data epochs. When we compared the adapted methods to the non-adapted ones on our simulated data, we generally did not find major differences.

As described for the AR-Fourier method, spectral estimation for the simulated data was performed at 4 Hz (the lower of the two simulated processes), and spectral estimation for the empirical data was performed at 10 Hz (the dominant frequency component identified by the FOOOF method).

#### 2.3.2. The Phase Locking Value as metric of phase estimation accuracy

We used the Phase Locking Value (PLV) as a measure of accuracy of spectral phase estimation, to compare the performance of various phase estimation methods. In our case, the computation of the PLV proceeded as follows: For each (empirically recorded or simulated) trial, the phase difference between the true phase and the estimated phase was calculated and expressed as unit-length complex vector. The complex vectors obtained across many trials were then averaged in the complex domain to obtain the residual vector. The length of this residual vector is the PLV. If the phase differences are random, the PLV tends towards zero for an infinite number of trials. If the phase differences are identical across trials, the PLV is one (Lachaux et al., 1999; Mormann et al., 2000). In our case, the closer the PLV was to one, the closer were the estimated phases to the true phases, and thus the better was the respective spectral estimation method. Note that in principle, the estimated phases could also have contained a consistent phase difference to the true phases, in which case the PLV would have merely measured the consistency of phase estimation, i.e., the degree to which phase estimation avoided noise, while it would not have measured any potential bias. However, we visually inspected the distributions of phase differences between estimated and true phases and found them, as expected, to be centered on zero.

Note that the AR-Fourier method entailed 100 extrapolations per trials. The spectral estimates of those 100 extrapolations were first averaged, and subsequently, the PLV was calculated over the trials. Thereby, the number of spectral estimates entering into the PLV calculation was identical between the AR-Fourier method and the other methods.

Note that the PLV contains a sample-size bias: If the true PLV is below one, the PLV estimate tends to the true PLV only for an infinite sample size, whereas for finite sample sizes, the PLV contains a systematic positive bias. This bias is small for the number of 500 trials used here. Most importantly, the number of trials was constant for all our analyses, such that differences in method performance are not due to differences in bias. For a true PLV of zero and 500 trials, the expected PLV is approximately 0.045. As we will show later, the estimated PLVs were highest for the AR-Fourier method and lower for the other methods. Thus, the PLVs for the other methods contained more of the positive sample-size bias, and this slightly reduced our estimates of the superiority of the AR-Fourier method.

#### 2.3.3. Comparison between methods and corresponding statistics

We compared the accuracy between the AR-Fourier method and alternative methods for simulated and empirical data.

For the simulated data, we were able to compare the AR-Fourier method to all seven alternative methods. We focused on a dataset with equal proportions of the 4 Hz and 6 Hz components, and with 90 combinations of pink-noise and phase-diffusion parameters. For each of the 90 parameter combinations, we simulated 500 trials. To compare the performance of AR-Fourier against the other seven methods, we used as test statistic the difference in PLV between AR-Fourier and each alternative method. The significance was assessed using a non-parametric randomization test, including correction for multiple comparisons. We performed 1000 randomizations. In each randomization, we performed the following steps: [1] For each of the 90 parameter combinations and for each of the seven methods combinations, [1.1] we randomly repartitioned the phase estimates obtained from that method and from AR-Fourier, respectively, keeping the number of phase estimates per method equal, [1.2] we calculated the surrogate PLVs and their differences between the alternative method and AR-Fourier; [2] We identified the maximum and minimum PLV difference across all parameter and method combinations, and placed the maximal difference into the max-randomization distribution, and the minimal difference into the min-randomization distributions. The 2.5^th^ percentile of the min-randomization distribution and the 97.5^th^ percentile of the max-randomization distribution were used as significance thresholds. This corresponds to a two-sided statistical test that controls the false-positive rate to be below 5% and corrects for the multiple comparisons across parameter and method combinations (Nichols and Holmes, 2002).

For the empirical data, we were able to compare the AR-Fourier method to six of the seven alternative methods. One of the alternative methods, the asymmetric-taper method referred to as Mitchell2007, would have required longer data segments than were available. In our comparison using empirical data, we utilized the Wilcoxon signed-rank test (as available in MATLAB) to compare paired samples of phase-locking values (PLVs) obtained from two different phase estimation methods. Specifically, we compared PLVs across channels, between the AR-Fourier method and each of the considered previously published methods. Correction for the multiple comparisons across the different combinations of methods was done with the Bonferroni method.

## 3. Results

### 3.1. Comparison to other methods

We compared our method against previously described methods, using both simulated and empirical data. First, we generated simulated data, for which the phase at the critical time is defined and thereby can be regarded as ‘true phase’. We then quantified the accuracy with which the true phase was estimated by calculating the PLV, across 500 simulated trials, between the true phases and the phase estimates from all considered methods. The simulations allow us to systematically manipulate the characteristics of the signal, specifically the degree of phase diffusion of the rhythmic component of the signal and the proportion of pink noise contained in the signal. By varying these simulation parameters over a wide range, we explored the influence of these factors on the quality of phase estimation. Furthermore, we explored whether differences in accuracy between methods generalize and thereby can be expected to occur similarly for a wide range of empirical data. Subsequently, we apply the different methods to empirical data recorded from awake macaque area V4, and again compare the quality of phase estimation.

#### 3.1.1. Simulated Data

We compared the accuracy of the AR-Fourier method to established techniques using simulated data created by combining two sinusoids of 4 Hz, the target component that we aimed to estimate, and 6 Hz, the distractor component, alongside 1/f noise, following a previously published comparison of spectral estimation methods (Wodeyar et al., 2021). To gauge method performance across datasets with diverse spectral characteristics, we manipulated various simulation parameters, including the phase diffusion of the signal, the intensity of 1/f noise, and the proportion of the 6 Hz component. Each phase-estimation method is then applied to these datasets with varying properties, as detailed in the methods section.

Overall, AR-Fourier provided the most accurate phase estimation for essentially all simulated datasets. The differences to alternative methods vanished primarily for phase-diffusion values approaching zero. A phase diffusion of zero corresponds to a simulated signal component that is sinusoidal, such that its phase evolves strictly linear with time and can therefore be estimated equally well for any part of the signal. Thus, for zero phase diffusion, spectral estimation at the edge is no different from general spectral estimation. Thus, the case of zero phase diffusion is only included here as sanity check and for completeness.

We found that the advantage of AR-Fourier decreased for decreasing strengths of the distractor component. Therefore, to keep the comparison conservative and to limit the number of variable parameters, we fixed the distractor component to be equally strong as the target component.

Figure 2 shows the accuracy of phase estimation, quantified as the PLV between the estimated phase and the ‘true phase’ of the simulated data, separately for each of the eight methods, as a function of both phase diffusion and noise coefficient of the simulated data. As expected, PLVs increased with decreasing noise coefficients and with decreasing phase diffusion.

**Figure 2.**
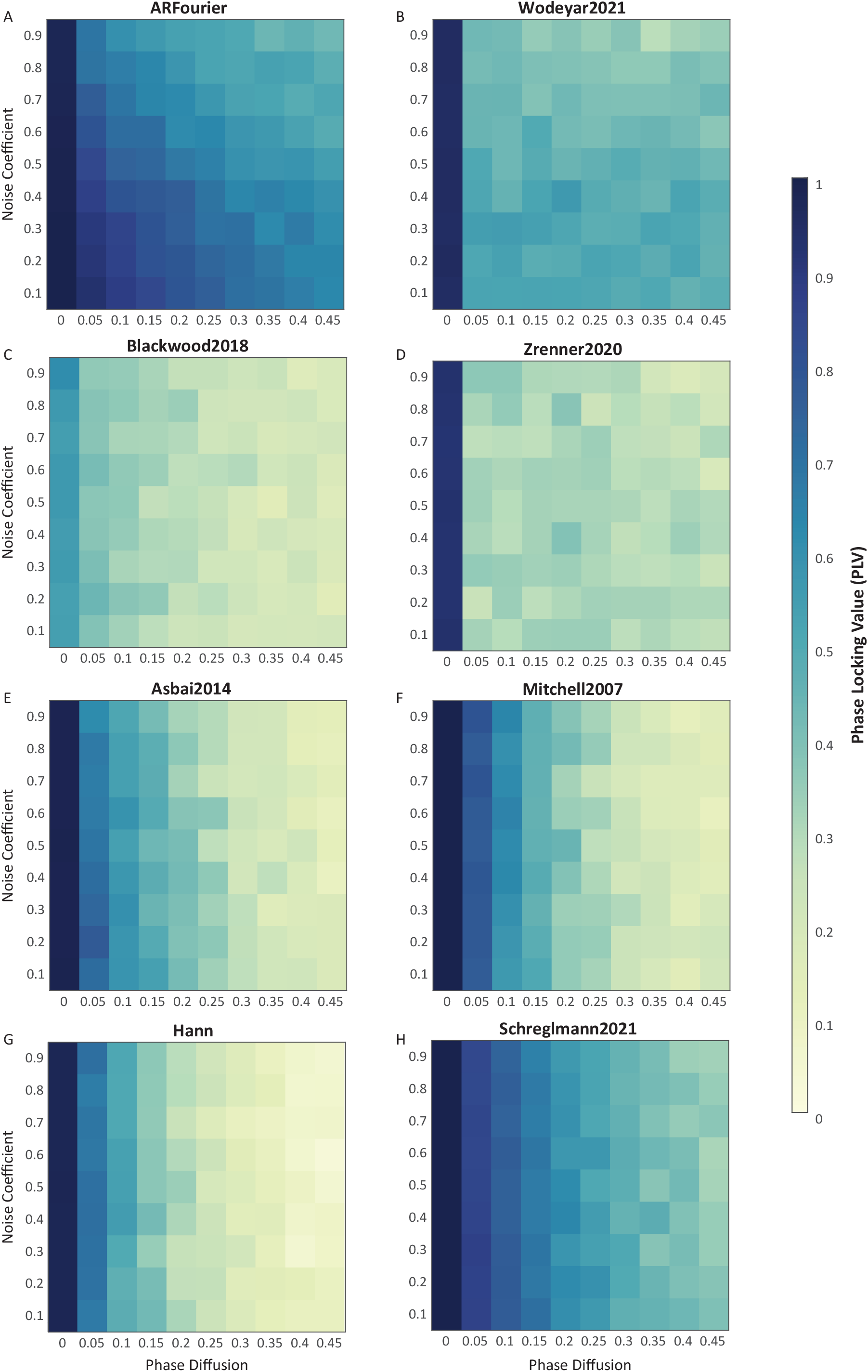
Accuracy of phase estimation across different methods for varying parameters of simulated data. Each subplot presents the phase estimation accuracy for each of the eight methods, including AR Fourier. The x-axis represents phase diffusion, and the y-axis represents noise coefficient. Each pixel indicates the phase locking value (PLV) between the estimated phases and the ‘true phases’, where values closer to 1 denote higher accuracy in phase estimation.

Figure 3 shows the comparison of AR-Fourier to each of the seven alternative methods. This is quantified by the difference in PLVs between AR-Fourier and each method, shown as a function of both phase diffusion and noise coefficient in the simulated data, similar to Figure 2. The figure also highlights the significant regions for these differences, indicating scenarios where AR-Fourier significantly outperforms or underperforms compared to the other methods. AR-Fourier shows a significantly better performance in most scenarios for most methods. A noteworthy exception is the Schreglmann2021 method, which showed a better performance than AR-Fourier for datasets with very low phase diffusion and very high noise.

**Figure 3.**
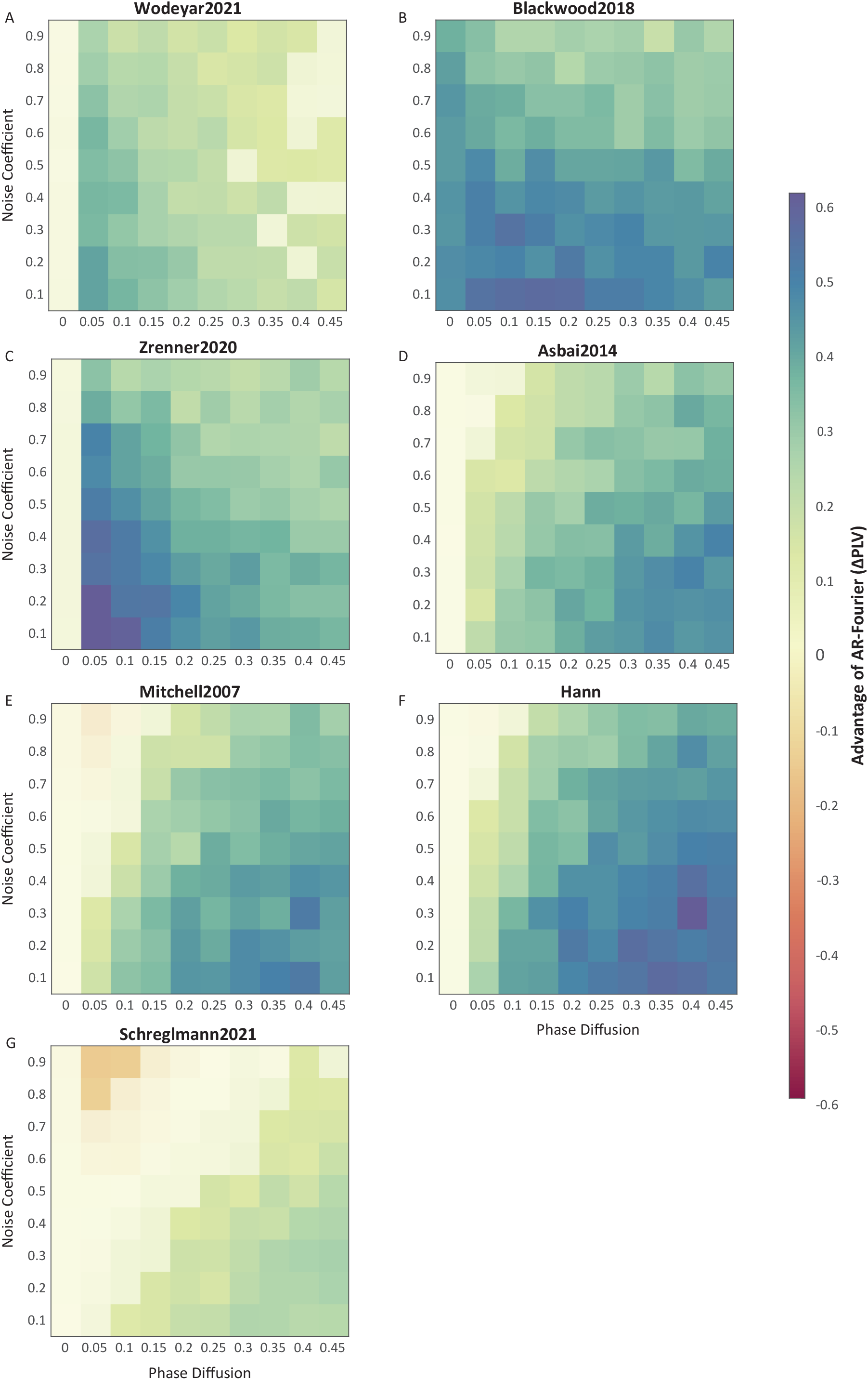
Comparison of AR-Fourier to alternative methods for varying parameters of simulated data. Each subplot presents the difference in phase estimation accuracy between AR-Fourier and each of the alternative methods. The x-axis represents phase diffusion, and the y-axis represents noise coefficient. Each pixel indicates the difference in PLVs between AR-Fourier and each of the methods. The pixels where the advantage (or disadvantage) of AR-Fourier is not significant are indicated by lower transparency.

Figure 4(A, B) shows cross sections through the 2D PLV matrices of Figure 2 at an intermediate noise coefficient of 0.5 (Figure. 4A), and at an intermediate phase diffusion of 0.25 (Figure. 4B). These cross sections suggest that the accuracy of phase estimation at the edge was particularly affected by phase diffusion. Figure 4(C, D) show the corresponding differences in accuracy between AR-Fourier and each alternative method. Figure 4C suggests that the advantage of AR-Fourier decreased slightly with increased phase diffusion compared to Zrenner2020 and Wodeyar2021, that it remained constant compared to Blackwood2018, and that it increased compared to Schreglmann2021 and the taper based methods Asbai2014, Mitchell2007 and Hann. Figure 4D suggests that the advantage of AR-Fourier decreased slightly with increasing noise coefficient compared to all alternative methods. Figure 4(C, D) also shows the significance thresholds, which reveals that the advantage of AR-Fourier was significant for most parameter combinations, except for the lowest phase-diffusion values. Note that the significance thresholds would be expected to shrink, rendering even smaller differences significant, for larger numbers of simulated trials, and we used 500 trials to be in the range often used in experiments.

**Figure 4.**
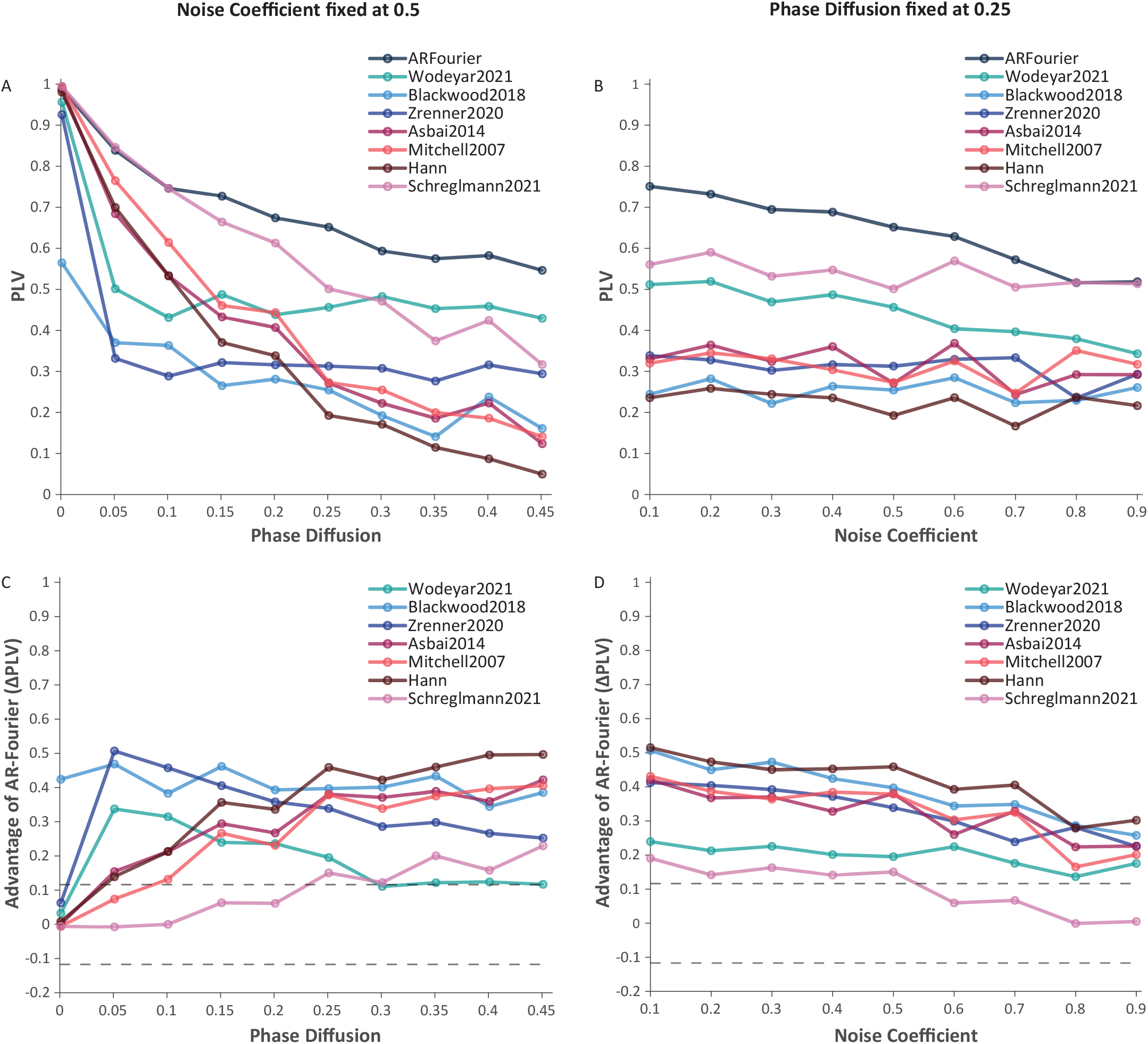
Accuracy and accuracy differences in cross sections through the simulated data. (A, B) Accuracy cross-sections at intermediate noise coefficient (0.5) and phase diffusion (0.25), each line represents PLV of a method. (C, D) Accuracy differences between AR-Fourier and each of the alternative methods at the fixed noise coefficient (0.5) and phase diffusion (0.25), with significance thresholds (dashed horizontal lines) to indicate AR-Fourier’s significant advantage (or disadvantage) over each of the alternative methods.

#### 3.1.2. Empirical Data

Having validated the accuracy of the AR-Fourier method across simulated data of variable characteristics, we next investigated its accuracy on a dataset recorded from a Utah array with 64 electrodes chronically implanted in awake macaque area V4. All electrodes, except one used as reference and a malfunctioning one, were used. Power spectra averaged over electrodes and trials revealed a dominant spectral component at 10 Hz, that is at the alpha rhythm, which we therefore used as the frequency for quantifying the accuracy of spectral estimation at the edge (see Methods for details).

Figure 5 shows scatter plots for each comparison of AR-Fourier to an alternative method. Each scatter plot shows 62 dots, corresponding to the 62 electrodes. The x- and y-values correspond to the respective PLVs across 500 trials, between a phase estimate at an edge versus a ‘true-phase’ estimate as described in the Methods. The advantage of AR-Fourier compared to all alternative methods was consistent across electrodes (Wilcoxon signed rank test, p = 4.5467e-11 after Bonferroni correction for six methods comparisons).

**Figure 5.**
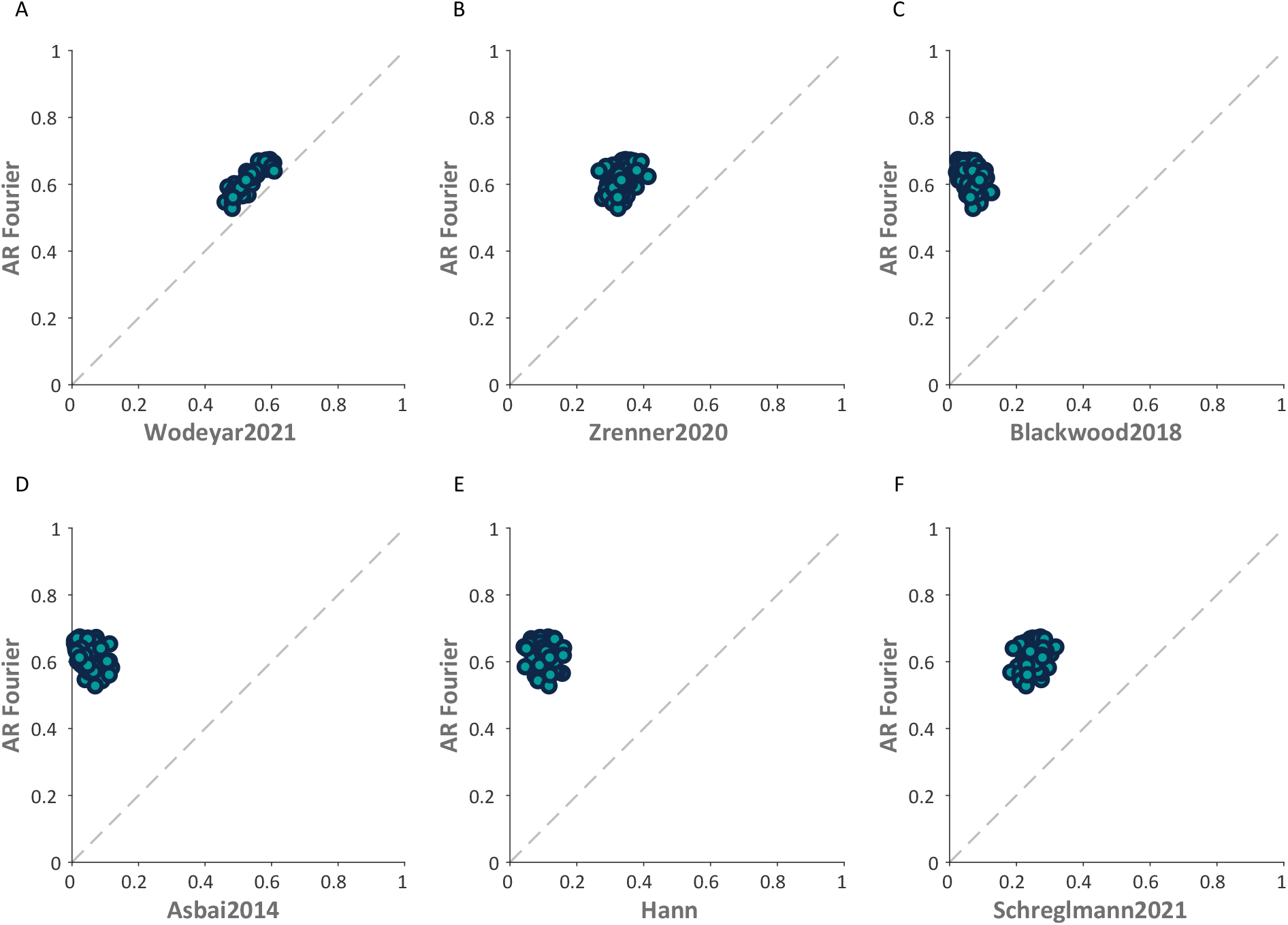
Accuracy of phase estimation across different methods for empirical data. Each subplot shows a scatter plot of phase locking values (PLVs) comparing AR-Fourier with each alternative method. Each dot represents one channel.

## 4. Discussion

We have developed a novel technique for estimating the spectral phase at the edge of a window, building on established extrapolation methods (Chen et al., 2013; Ni et al., 2016; Blackwood et al., 2018; Zrenner et al., 2020; Wodeyar et al., 2021; Wodeyar et al., 2023). This new approach aims at overcoming some limitations associated with existing extrapolation-based methods, particularly those arising from the challenges of band-pass filtering data. Band-pass filtering can introduce significant group delays that vary with frequency, potentially shifting the critical time in a frequency-dependent manner and complicating analysis. To address this issue, we propose an autoregressive (AR) extrapolation-based method that circumvents band-pass filtering by utilizing the broadband signal. Additionally, our approach includes a technique to average across multiple extrapolations, enhancing precision and reliability in phase estimation.

We evaluated the effectiveness of our AR-Fourier method through testing against established methods using simulated and empirical datasets. In the simulated data, we varied parameters such as phase diffusion, the proportion of 1/f noise, and the proportion of the distractor component. Our findings revealed that while some methods exhibit comparable performance to AR-Fourier under conditions of low phase diffusion and noise, they struggle under high phase diffusion and noise levels. While AR-Fourier also suffered from increasing phase diffusion and/or noise levels, it did so to a lesser degree than the other methods. This is demonstrated by the significant advantage in the PLVs of AR-Fourier compared to each of the other methods for most combinations of phase diffusion and noise.

When considering all combinations of phase diffusion and noise in the simulated datasets, Schreglmann2021 showed better performance than AR-Fourier in scenarios with very low phase diffusion and very high noise (Figure. 3G). We can speculate on the reasons for this. When noise is high, it should be averaged out by the spectral estimation process integrating over long estimation windows. Such long estimation windows can be afforded when phase diffusion is low. Scenarios of low phase diffusion and high noise should therefore be well handled by approaches that integrate spectral information over long time windows. A very long time window of 5 s was provided to the Schreglmann2021 method for the simulated data, and this is where we observed outperformance for the specific parameters. However, in our experience, 5 s of (relatively) stationary data before a critical time is typically not available in empirical datasets. Indeed, the empirical dataset used here provided 0.5 s of data before the critical time, and for this empirical dataset, the Schreglmann2021 method gave less accurate estimates than the AR-Fourier method (Figure. 5F). Note that specific estimation window lengths might be advantageous for specific combinations of noise and phase diffusion. This also entails that any one of the methods can most likely be optimized for specific datasets by tuning their free parameters, like e.g. the specific shape of the asymmetric tapers, the width of the frequency-domain filter for Schreglmann2021 or the number of cycles for the AR-Fourier method.

Real-world datasets might exhibit high levels of phase diffusion, noise, and multiple concurrent frequencies. Yet, the specific combination of phase diffusion and noise in empirical data is not known. Therefore, we aimed at comparing the quality of spectral estimation at the edge in empirical data. We used LFP data from a Utah electrode array chronically implanted in area V4 of a macaque monkey, recorded while the animal was awake and performing an attentional task. In this dataset, we observed significant phase estimation accuracy improvements with AR-Fourier compared to previously described methods. A visual inspection of Figure 5 suggests that for those empirical data, the closest contender to AR-Fourier was the Wodeyar2021 method. Surprisingly, the Schreglmann2021 method, which outperformed AR-Fourier for simulated data with very low phase diffusion and very high noise, performed relatively poorly on this empirical dataset. It is noteworthy that in the in vivo signal, the intensity of rhythms and the level of phase diffusion and noise most likely change with time. Moreover, these parameters most likely vary across channels. Nonetheless, the AR-Fourier method demonstrated a strong performance on this dataset, indicating its practical utility for empirical data analysis. These results underscore the robustness of AR-Fourier, making it a promising choice for applications demanding precise phase estimation in complex signals.

Despite its advantages, it is essential to acknowledge certain limitations of our method. One notable challenge is its computational complexity and time-consuming nature, especially when conducting multiple extrapolations. This factor should be taken into account when selecting a phase estimation method in real-time or resource-limited scenarios.

When choosing phase estimation methods, accuracy is an important but not the only consideration as implied above. It is essential to consider the nature of the data, available resources and the specific requirements for the analysis. For instance, the parameter of phase diffusion plays a critical role in method selection. Understanding the decorrelation time of the signal, which is linked to phase diffusion, can offer valuable insights into the data’s characteristics and guide method selection. The decorrelation time of a signal can be approximated with the following procedure: Start by applying a band-pass filter to isolate the major frequency range of interest. Then, calculate the autocorrelation function of the filtered data and fit an exponential curve to it. The decorrelation time can then be approximated as the reciprocal of the time constant obtained from this fit. The decorrelation time can be mapped to a phase diffusion value using our approach of simulating datasets with different phase diffusion values. This process provides valuable information about the temporal characteristics of the signal, which is particularly useful when dealing with significant phase diffusion, ultimately assisting in selecting appropriate methods for data analysis.

In scenarios with high phase diffusion (i.e. short decorrelation time), we recommend using either the AR-Fourier or Wodeyar2021 methods based on available computational and time resources. If computational resources are limited and the analysis does not demand high precision, the Wodeyar2021 method is suitable. Otherwise, opt for the AR-Fourier method to ensure greater accuracy and reliability in phase estimation. Moreover, in scenarios with low phase diffusion and high noise, asymmetric tapers or the Schreglmann2021 approach might be more suitable for phase estimation. By considering these factors alongside accuracy, researchers can ensure that the selected method aligns with the specific requirements and properties of their dataset, leading to more effective analysis.

The significance of improving phase estimation methods extends beyond theoretical implications, with practical implications for understanding brain dynamics and advancing therapeutic interventions. Brain rhythms’ phase has been demonstrated to have an impact on cognitive function in various studies. It has been suggested that phase plays a significant role in perception (Busch et al., 2009; VanRullen, 2016; Fiebelkorn et al., 2018) and is implicated in communication at both local and global scales (Roberts et al., 2013; Fries, 2015; Maris et al., 2016). Additionally, studies indicate that phase is involved in coordinating neuronal spikes and local field potentials (Ray et al., 2008; Zanos et al., 2012; Pesaran et al., 2018). Phase-amplitude synchrony, commonly observed in neural activity, may be altered in neurodegenerative diseases, suggesting potential implications for disease detection (de Hemptinne et al., 2013). Moreover, phase holds significance in therapeutic interventions (Ngo et al., 2013; Cagnan et al., 2017).

The estimation methods considered here are also applicable in real-time phase determination for closed-loop brain stimulation. Studies on therapeutic interventions propose that phase-based stimulation can be effective in mitigating symptoms of Parkinson’s disease (Cagnan et al., 2017; Zanos et al., 2018). For instance, the Zanos et al. (2018) study employed brain stimulation based on phase information derived from immediately preceding data, requiring precise phase estimation at the signal epoch’s edge. Achieving such precision at the edge necessitates the development of sophisticated estimation methods like the one proposed in this paper. The AR-Fourier method represents a notable advancement in phase estimation methods, offering enhanced accuracy and reliability for complex real-world signals. Future research should focus on optimizing computational efficiency and integrating our method into practical applications, such as closed-loop brain stimulation systems.

In summary, improving phase estimation methods not only enhances our understanding of brain dynamics but also holds promise for advancing therapeutic interventions, particularly in real-time applications such as closed-loop brain stimulation. The development and refinement of such methods contributes to the ongoing progress in neuroscientific research and clinical applications.

## Data Availability

The data and method implementations will be made available upon publication.

## Author Contribution

Shivangi Patel: Conceptualization, Formal analysis, Methodology, Software, Visualization, Writing - Original Draft, Writing - Review & Editing. Elena Psarou: Conceptualization, Investigation, Methodology, Supervision, Writing - Review & Editing. Gregor Mönke: Methodology, Software, Writing - Review & Editing. Pascal Fries: Conceptualization, Funding acquisition, Project administration, Resources, Supervision, Writing - Original Draft, Writing - Review & Editing.

## Acknowledgements

We thank Tim Näher for helpful discussions and suggestions.

## Funding Information

P.F. was supported by: the German Research Foundation (DFG, FR2557/2-1, FR2557/5-1, FR2557/7-1), the European Union (FP7-604102-HBP), the National Institutes of Health (NIH, 1U54MH091657-WU-Minn-Consortium-HCP).

## Declaration of Interests

P.F. has a patent on thin-film electrodes and is member of the Advisory Board of CorTec GmbH (Freiburg, Germany). The other authors declare to have no competing interests.

